# Genetic epidemiology and Mendelian randomization for informing disease therapeutics: conceptual and methodological challenges

**DOI:** 10.1101/126599

**Authors:** Lavinia Paternoster, Kate Tilling, George Davey Smith

## Abstract

The past decade has been proclaimed as a hugely successful era of gene discovery through the high yields of many genome-wide association studies (GWAS). However, much of the perceived benefit of such discoveries lies in the promise that the identification of genes that influence disease would directly translate into the identification of potential therapeutic targets (1-4), but this has yet to be realised at a level reflecting expectation. One reason for this, we suggest, is that GWAS to date have generally not focused on phenotypes that directly relate to the progression of disease, and thus speak to disease treatment.

---

As of 2017-04-03, the GWAS Catalog contained 2854 publications and 33674 unique SNP-trait associations (5). The large majority of these studies investigate genetic variation related to the presence (or occurrence) of disease. Such variants, though they may be informative for prevention of disease, have unclear utility in informing disease treatment. If variants implicate aetiological mechanisms of importance for disease onset, but of little relevance to disease progression, then the use of case/control GWAS as evidence to inform disease treatment related drug discovery could be futile. As an obvious example consider GWAS of lung cancer. The lead variants identified in such GWAS tag a locus related to heaviness of cigarette smoking (6), supporting the overwhelming evidence that smoking causes lung cancer. However, cessation of smoking is hardly an efficacious treatment strategy after the onset of disease, although not smoking is a highly effective means of very substantially reducing the risk of developing lung cancer in the first place. Examples of factors causing both disease incidence and disease progression exist - for example, LDL cholesterol levels clearly influence risk of initial coronary events and lowering LDL cholesterol reduces risk of subsequent events. However, it is not necessarily the case that risk factors will influence both disease onset and disease progression – for example, a recent GWAS of Crohn’s disease observed independent genetic variants for risk of onset and progression (7), and reported a negative genetic correlation (estimated through LD score regression) between occurrence and progression, although this was imprecisely estimated. It is indeed possible that in some cases the effects of a particular exposure on initiation and prognosis of disease could be in opposite directions, as has been suggested with respect to folate intake and colon cancer(8).

In contrast to the large body of research on genetic risk of disease incidence, only a small proportion of GWAS studies (∼8% of associations curated in the GWAS Catalog (p<1 × 10 ^-5^)) have attempted to identify variants associated with disease progression or severity, and those that have are mostly small (90% have n<5000). Investigating disease progression as a trait offers considerable opportunity for identifying treatment targets and informing therapeutics, but it also introduces several important complications that have had little formal discussion in the literature and have not been addressed in many of the existing disease progression studies. A key problem, which we will discuss in more detail, is the issue of potential introduction of collider bias when studying a selected (i.e. case-only) group of individuals.

GWAS studies are now also routinely being used to help strengthen causal inference with respect to observational associations between exposures and disease, using Mendelian Randomization (MR) (9, 10), (see BOX 1). With its emphasis on causality it is important to appreciate that the challenges we present here also apply to MR. To date, few studies have used MR to identify factors influencing disease progression. In the supplementary table 1 we summarise the 27 MR studies of progression that we identified in a systematic search. Only one of these studies (9) acknowledged the issue of potential introduction of confounding through collider bias; interestingly this was the first of these studies to be published.

**Table 1:** Estimated effects of A (risk factor for incidence only) and C (risk factor for incidence and progression) from Figure 1, under different degrees of unmeasured confounding of incidence and progression.

|  | Degree of confounding by unmeasured confounder(s) (U) |  |  |  |
| --- | --- | --- | --- | --- |
|  | Low | Mod | High | Strong |
|  | OR for disease=1.5<br>Beta for progression=0.5 | OR for disease=2<br>Beta for progression=0.8 | OR for disease=2.5<br>Beta for progression=1 | OR for disease=3<br>Beta for progression=1.5 |
| **Apparent effect of A on progression**<br>(regression coefficient, SE)<br>**True effect=0** | -0.01 (0.01) | -0.02 (0.02) | -0.03 (0.02) | -0.06 (0.03) |
| Percentage of 95% CI including 0 | 90% | 78% | 66% | 35% |
| **Apparent effect of C on progression**<br>(regression coefficient, SE)<br>**True effect=0.1** | 0.10 (0.01) | 0.08 (0.01) | 0.07 (0.01) | 0.04 (0.02) |
| Proportion of 95% CI including 0.1 | 72% | 35% | 18% | 1% |

Each cell represents results from 500 simulations with a sample size of 50,000.
Uppercase letters refer to factors in Figure 1, lowercase letters refer to effect sizes of paths in Figure 1.
In all scenarios the OR for A and C for disease incidence are 1.3, and the MAF for genetic variants A is 0.2.
C and the unmeasured confounder (U) are standard normal variables, disease is a binary variable (with prevalence of approximately 0.2) and prognosis is a normally distributed variable.

### BOX 1: Mendelian Randomization

Mendelian randomization is an approach that uses genetic variation to improve causal inference in observational studies. A genetic variant associated with the exposure of interest (genetic instrument) is used to test the causal relationship between exposure and outcome (Figure B1). If there is association between the genetic instrument and the outcome, then there is assumed to be a causal relationship, because unlike in the observational association, the genetic variant is not subject to issues of reverse causation and/or confounding.

**Figure B1.**
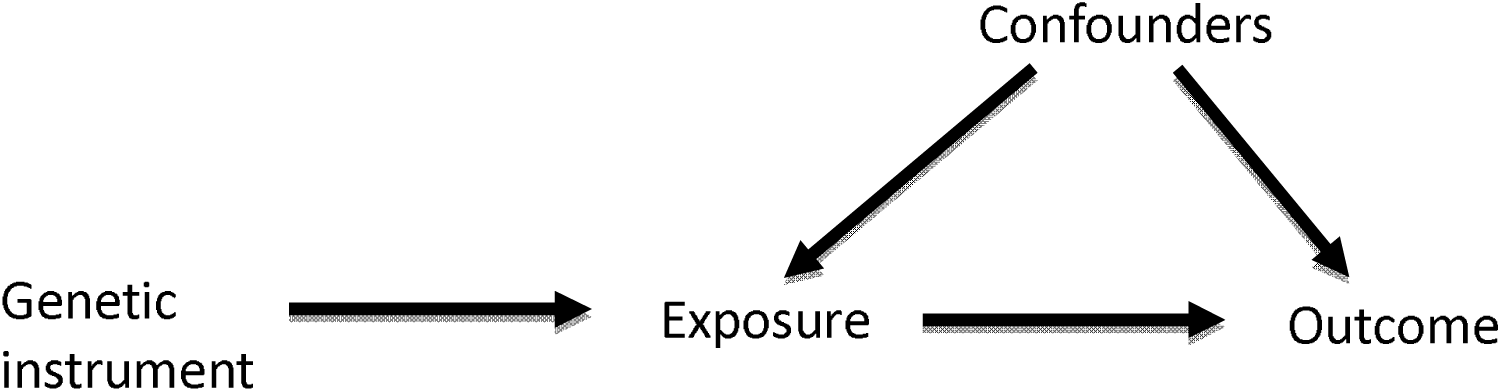
Directed acyclic graph (DAG) of Mendelian Randomization method

Assumptions of MR (19):

1. The genetic instrument is associated with the exposure of interest
2. The genetic instrument is independent of factors that confound the association of the exposure and the outcome
3. The genetic instrument is independent of the outcome, given the exposure and the confounders

The method has been widely applied in the investigation of exposures that increase the risk of disease (20), both within single studies and in a two-sample framework based on summary data, generally from large-scale genome wide association study (GWAS) consortia (21). Such studies have demonstrated evidence of causal relationships (e.g. for obesity, blood pressure and smoking with increased risk of coronary heart disease CHD (22-24)), lack of causal relationships (e.g. for C reactive protein relationship with CHD, diabetes and cancer (25-27)), debunking supposed protective behaviours (such as the beneficial effects of moderate alcohol intake on CHD risk (28)) and predicting randomised controlled trial successes and failure (29).

The emphasis on causality in a Mendelian randomization study has led to the acknowledgement within the field that they are also likely to have great value in suggesting what are likely to be successful interventions for treatment of disease (30,31). However, there are particular aspects of the study of disease prognosis that limit the applicability of Mendelian randomization.

## Challenges for genetic and MR studies of disease progression

### Collider bias

Collider bias is a fundamental issue in progression studies(11) (Figure 1). When a study group are selected on certain characteristics (e.g. being cases for a particular disease), this will introduce inverse associations between all independent risk factors for characteristics relating to being included within the study sample. For example, in a study of CHD progression, where only CHD cases are selected for inclusion, there will be associations induced between all CHD risk factors (genetic and non-genetic) amongst the study individuals. Therefore, in a genetic study of progression within these cases, collider bias will induce spurious associations between genetic variants and progression (providing that at least one other factor influences both incidence and progression) (12). Similarly, in an MR study of progression within these cases, the assumption that ‘the genetic instrument is independent of factors that confound the association of the exposure and the outcome’ (assumption 2, BOX 2) would be violated.

**Figure 1.**
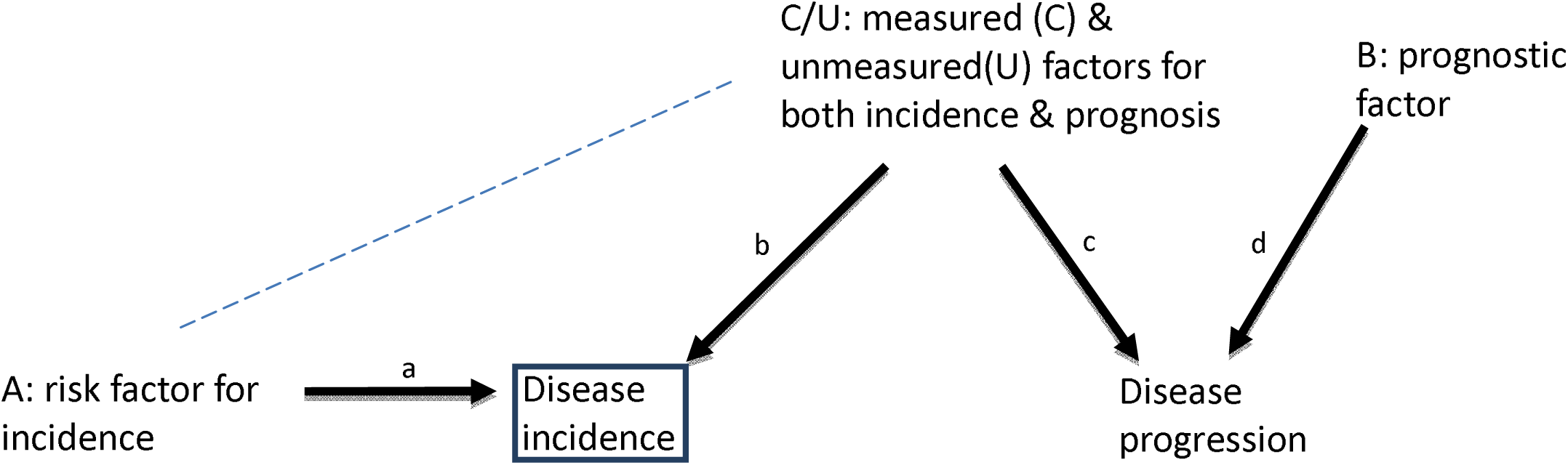
Directed acyclic graph (DAG) demonstrating the issue of collider bias in studies with participants selected according to disease status. In this situation collider bias induces an inverse association (dashed line) between any factors (A, C and U) that affect disease incidence (or other study selection criteria). When one or more of these factors are also associated with disease progression (C, U), a back-door path is opened up from A to disease progression through the induced association. If A is a genetic risk factor, it can appear there is an association between genetic risk factor A and disease progression only because of the induced association with C or U. If C is measured and can be adjusted for, the induced association is blocked, but unmeasured U cannot be adjusted for in the analysis. Only when the genetic risk factor for progression is not also a risk factor for incidence (i.e. B) will it not be affected by selection bias.

##### BOX 2: Mendelian Randomization

Collider bias is an issue in MR of progression, because for any exposure that causes onset of disease, the genetic instruments for that exposure will be inversely associated with any other risk factor for onset and so the association between the genetic variant and progression may be subject to confounding by these factors (Figure B2). Although this is true for single variants, the combination of variants into a polygenic score may serve to dramatically increase this effect (32).

**Figure B2.**
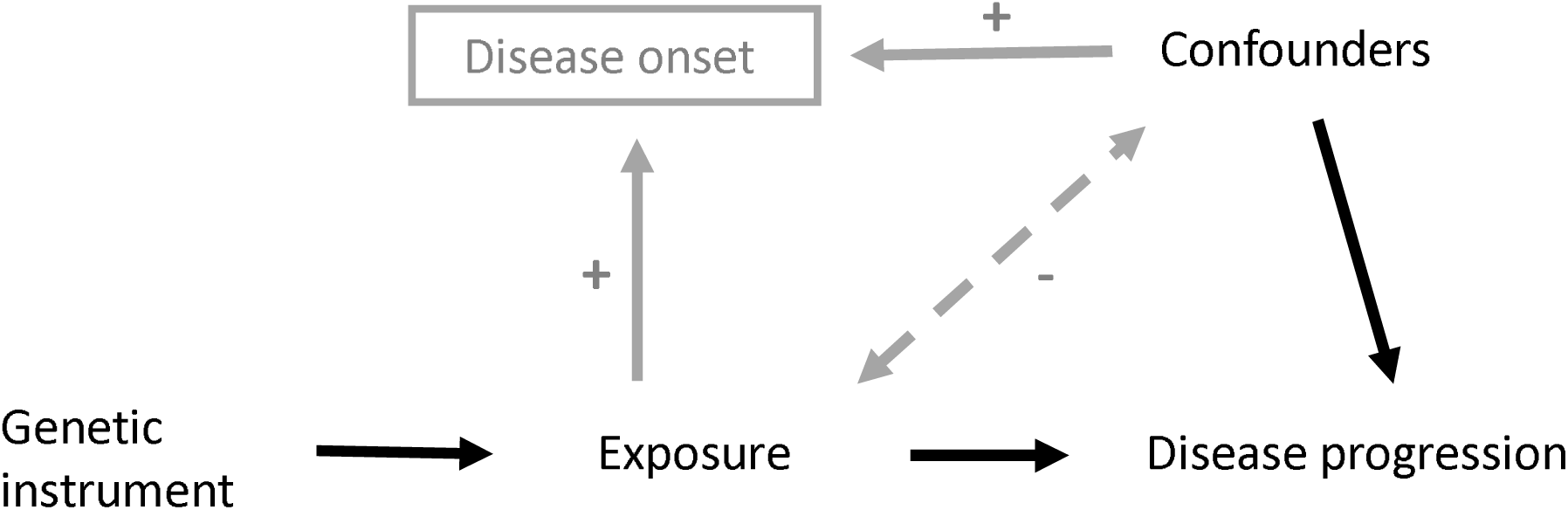
DAG to demonstrate how the introduction of collider bias through the selection of cases (grey paths) can impact an MR analysis between an exposure and disease progression as an outcome.

Association induced because SNP causes disease (via exposure), and thus conditioning on disease induces an association between all variables causing disease. In a model not adjusting for exposure (e.g. relating progression to SNP), there is an association between SNP and the confounders, which biases the SNP-progression association.

We investigated the bias due to studying cases only using a simple simulation study (Table 1). We simulated the situation depicted in Figure 1, with both a measured (C) and an unmeasured (U) confounder of disease incidence and progression. We simulated situations with low, moderate, high and strong confounding. Collider bias has somewhat different implications for two underlying biological mechanisms. One (as depicted in Figure 1), where risk factor A causes disease incidence, but A does not cause disease progression. In this scenario, studying cases only introduces collider bias, which induces an association between A and C, and thus results in an induced association between A and disease progression in the study sample (Table 1). The bias in the estimated effect of A on disease progression increases as the degree of unmeasured confounding of disease incidence and progression increases (i.e. the degree to which there are common factors which influence disease onset and progression), with the proportion of 95% confidence intervals including the true effect of zero falling from 90% (low confounding) to 35% (strongest confounding). The second scenario, is where risk factor C causes both disease incidence and progression (Figure 1). Collider bias is again induced by studying only cases, and here it biases the estimated effect of C on progression towards the null (Table 1). Again, the bias increases as the degree of confounding of incidence and progression increases.

This collider bias can lead to either over- or under-identification of genetic risk factors for progression, depending on the direction of the relationships between the risk factors and disease onset. Collider bias should always be properly considered and a number of things can be done to mitigate this potential bias.

1. Check for association between the genetic variant and disease incidence in any study of disease progression. When a variant is identified as associated with progression, the association between this variant and disease incidence (or other selection criteria) should also be reported. This can demonstrate whether there is any potential for collider bias.
2. Check for associations between the genetic variant and potential confounders in the study sample – such associations might indicate that both the genetic variant and confounders influence disease incidence(13).
3. If there are associations between genetic variant and potential confounders of disease incidence and progression, then adjusting for such confounders will mitigate the problem. However, investigators should be aware that as with any study of traditional risk factors, unmeasured confounding will remain an issue.
4. If certain parameters are known (such as prevalence of disease and the effects of the genetic and potential confounders on disease onset), then it is possible to estimate the induced bias and so potentially correct for it using analytical formulae (12) or inverse probability weighting.

It is an important aside to note that whilst disease incidence and diagnosis are the particular selection criteria of concern in the context of a progression study, ANY factor which relates to selection of study participants can result in collider bias(11). Therefore, any study where the participants are not a random selection of the population can suffer from induced association between genetic variants and factors which are independent in the underlying population.

### Confounding with disease stage at baseline

Studies of progression should be carefully designed so that it is true ‘progression’ that is the outcome. Under some situations disease detection (and hence position of individuals along the disease progression timeline at diagnosis) may be associated with other factors (e.g. smoking could be related to age at onset). For example, suppose that older people were more likely to take part in a screening programme, as national screening programmes often have a lower age limit. Thus, older people with cancer would tend to have their cancer detected earlier (by screening), and thus present with less advanced cancer, whereas younger people with cancer might present with symptomatic (more advanced) cancer. In a study of people with this cancer, it would appear that age was a positive prognostic factor. However, if stage at study entry was assessed, then the association between age and stage could be examined, and controlled for in the analysis. Ideally stage of disease at study entry should be independent of the genetic variants. Collider bias with factors such as age might violate this – if age and genetic variant both influence disease incidence, and age influences stage of disease at study entry, then in a case-only study, the genetic variant would appear to be associated with age and hence, also with stage of disease at study entry. In this example, this spurious correlation could be removed by adjusting for age – however, in practice, all the factors influencing risk of disease occurrence will not be known.

### Measurement of progression

GWAS and MR typically use a single measure of either a continuous (e.g. blood pressure at age 60) or a binary (e.g. occurrence of a myocardial infarction by age 60) outcome. In a study of progression, the outcome may be more complex: time to cancer recurrence; survival time; accumulation of disability over a 20-year period; or recurrence-free survival time. For these outcomes more sophisticated analysis may be required such as survival analysis (including handling censoring - whereby follow-up data may be missing for individuals in a non-random pattern) and analysis of trajectories. We have developed methodology for GWAS of trajectories(14, 15), and methods for Mendelian Randomization in the context of survival analysis are available(16) but computational challenges remain and further methodological development is much needed. In addition, to allow well-powered meta-analysis studies to be conducted, comparable measures of progression will need to be available across datasets.

### Availability of data

GWAS and MR of disease occurrence has had huge recent success, in no small part due to the availability of very large datasets. In order for GWAS and MR of progression to see the same success, there is a need for availability of large-scale studies with both progression and genetic data. One potential source of such data is from randomised controlled trials, which will have detailed follow-up of patients and often now collect DNA as a standard. Genome-wide genotyping of such resources is an important first step. Generation of valuable progression data for GWAS is likely to require large consortia collaboration (as has been the case for traditional GWAS). Therefore, standardisation of progression measures across a number of studies is also going to be important for this approach to reach its full potential.

If all of these issues are appropriately addressed, there is huge opportunity for GWAS and MR of disease progression to identify potential new treatments(17). Platforms such as MR-Base (18), which catalogues all available GWAS data for simple implementation of MR, will make it possible to easily screen for potential new drug targets to treat disease.

## Acknowledgments

The authors work in an MRC-funded unit (MC_UU_12013/1, /4, /9)

## References

1. Cowan WM, Kopnisky KL, Hyman SE. The human genome project and its impact on psychiatry. Annu Rev Neurosci. 2002;25:1–50.

2. Obeidat M, Hall IP. Genetics of complex respiratory diseases: implications for pathophysiology and pharmacology studies. British journal of pharmacology. 2011;163(1):96–105.

3. Manolio TA, Collins FS. The HapMap and genome-wide association studies in diagnosis and therapy. Annual review of medicine. 2009;60:443–56.

4. Gabbani T, Deiana S, Marocchi M, Annese V. Genetic risk variants as therapeutic targets for Crohn’s disease. Expert opinion on therapeutic targets. 2017;21(4):381–90.

5. www.ebi.ac.uk/gwas/.

6. Munafo MR, Timofeeva MN, Morris RW, Prieto-Merino D, Sattar N, Brennan P, et al. Association between genetic variants on chromosome 15q25 locus and objective measures of tobacco exposure. Journal of the National Cancer Institute. 2012;104(10):740–8.

7. Lee JC, Biasci D, Roberts R, Gearry RB, Mansfield JC, Ahmad T, et al. Genome-wide association study identifies distinct genetic contributions to prognosis and susceptibility in Crohn’s disease. Nat Genet. 2017;49(2):262–8.

8. Kim YI. Role of folate in colon cancer development and progression. J Nutr. 2003;133(11 Suppl 1):3731S–9S.

9. Smith GD, Ebrahim S. ‘Mendelian randomization’: can genetic epidemiology contribute to understanding environmental determinants of disease? Int J Epidemiol. 2003;32(1):1–22.

10. Davey Smith G, Hemani G. Mendelian randomization: genetic anchors for causal inference in epidemiological studies. Human molecular genetics. 2014;23(R1):R89–98.

11. Munafo MR, Tilling K, Taylor AE, D.M. E, Davey Smith G. Collider Scope: How selection bias can induce spurious associations. BioRxiv http://doiorg/101101/079707. 2017.

12. Yaghootkar H, Bancks MP, Jones SE, McDaid A, Beaumont R, Donnelly L, et al. Quantifying the extent to which index event biases influence large genetic association studies. Human molecular genetics. 2017;26(5):1018–30.

13. Balazard F, Le Fur S, Bougnères P, Valleron A-J, group tI-Dc. Disease as collider: a new case-only method to discover environmental factors in complex diseases with genetic risk estimation. BioRxiv http://doiorg/101101/124560. 2017.

14. Warrington NM, Howe LD, Paternoster L, Kaakinen M, Herrala S, Huikari V, et al. A genome-wide association study of body mass index across early life and childhood. International journal of epidemiology. 2015;44(2):700–12.

15. Warrington NM, Tilling K, Howe LD, Paternoster L, Pennell CE, Wu YY, et al. Robustness of the linear mixed effects model to error distribution assumptions and the consequences for genome-wide association studies. Statistical applications in genetics and molecular biology. 2014;13(5):567–87.

16. Tchetgen Tchetgen EJ, Walter S, Vansteelandt S, Martinussen T, Glymour M. Instrumental variable estimation in a survival context. Epidemiology (Cambridge, Mass). 2015;26(3):402–10.

17. Davey Smith G, Paternoster L, Relton C. When Will Mendelian Randomization Become Relevant for Clinical Practice and Public Health? Jama. 2017;317(6):589–91.

18. Hemani G, Zheng J, Wade K, Laurin C, Elsworth B, Burgess S, et al. MR-Base: a platform for systematic causal inference across the phenome using billions of genetic associations. BioRxiv https://doi.org/10.1101/0789722016.

19. Lawlor DA, Harbord RM, Sterne JA, Timpson N, Davey Smith G. Mendelian randomization: using genes as instruments for making causal inferences in epidemiology. Statistics in medicine. 2008;27(8):1133–63.

20. Boef AG, Dekkers OM, le Cessie S. Mendelian randomization studies: a review of the approaches used and the quality of reporting. Int J Epidemiol. 2015;44(2):496–511.

21. Pierce BL, Burgess S. Efficient design for Mendelian randomization studies: subsample and 2-sample instrumental variable estimators. Am J Epidemiol. 2013;178(7):1177–84.

22. Emdin CA, Khera AV, Natarajan P, Klarin D, Zekavat SM, Hsiao AJ, et al. Genetic Association of Waist-to-Hip Ratio With Cardiometabolic Traits, Type 2 Diabetes, and Coronary Heart Disease. Jama. 2017;317(6):626–34.

23. Varbo A, Benn M, Davey Smith G, Timpson NJ, Tybjaerg-Hansen A, Nordestgaard BG. Remnant cholesterol, low-density lipoprotein cholesterol, and blood pressure as mediators from obesity to ischemic heart disease. Circulation research. 2015;116(4):665–73.

24. Asvold BO, Bjorngaard JH, Carslake D, Gabrielsen ME, Skorpen F, Davey Smith G, et al. Causal associations of tobacco smoking with cardiovascular risk factors: a Mendelian randomization analysis of the HUNT Study in Norway. International journal of epidemiology. 2014;43(5):1458–70.

25. C Reactive Protein Coronary Heart Disease Genetics Collaboration, Wensley F, Gao P, Burgess S, Kaptoge S, Di Angelantonio E, et al. Association between C reactive protein and coronary heart disease: mendelian randomisation analysis based on individual participant data. BMJ. 2011;342:d548.

26. Brunner EJ, Kivimaki M, Witte DR, Lawlor DA, Davey Smith G, Cooper JA, et al. Inflammation, insulin resistance, and diabetes-Mendelian randomization using CRP haplotypes points upstream. PLoS medicine. 2008;5(8):e155.

27. Allin KH, Nordestgaard BG, Zacho J, Tybjaerg-Hansen A, Bojesen SE. C-reactive protein and the risk of cancer: a mendelian randomization study. Journal of the National Cancer Institute. 2010;102(3):202–6.

28. Holmes MV, Dale CE, Zuccolo L, Silverwood RJ, Guo Y, Ye Z, et al. Association between alcohol and cardiovascular disease: Mendelian randomisation analysis based on individual participant data. Bmj. 2014;349:g4164.

29. Stitziel NO, Kathiresan S. Leveraging human genetics to guide drug target discovery. Trends in cardiovascular medicine. 2016;S1050-1738(16):30128–1.

30. Ference BA, Robinson JG, Brook RD, Catapano AL, Chapman MJ, Neff DR, et al. Variation in PCSK9 and HMGCR and Risk of Cardiovascular Disease and Diabetes. N Engl J Med. 2016;375(22):2144–53.

31. Mokry LE, Ahmad O, Forgetta V, Thanassoulis G, Richards JB. Mendelian randomisation applied to drug development in cardiovascular disease: a review. Journal of medical genetics. 2015;52(2):71–9.

32. Martin J, Tilling K, Hubbard L, Stergiakouli E, Thapar A, Davey Smith G, et al. Association of Genetic Risk for Schizophrenia With Nonparticipation Over Time in a Population-Based Cohort Study. Am J Epidemiol. 2016;183(12):1149–58.

